# Text Mining miRNA-gene Interactions for the mx-plore Platform

**DOI:** 10.1101/2025.01.22.634367

**Authors:** Markus Joppich, Samuel Klein, Ralf Zimmer

**Affiliations:** Ludwig-Maximilians-Universtät München, Department of Informatics, Institute of Bioinformatics, Munich, Germany

## Abstract

miRNAs are post-transcriptional regulators which bind to the specific mRNA of expressed genes and induce increased mRNA degradation, thereby reducing the gene expression of their target genes. miRNAs perform their regulatory functions in a context-specific manner. Several databases contain collections of miRNA-target interactions, which are either experimentally validated, derived from high-throughput experiments, or computationally predicted or captured from existing literature. For most databases, the specific context of the miRNA-target interaction is unknown or missing in these databases. mx-plore is not only a database of text-dervied miRNA-gene interactions, but it enhances the found interactions including detailed context information from scientific publications. To derive the entries in the mx-plore database, a newly developed text mining strategy combining dependency graph analysis and rule-based systems has been used. The mx-plore platform makes miRNA-gene interactions accessible and searchable on various levels, such as by cell type, disease or involved processes. The platform is available at https://rest.bio.ifi.lmu.de/mxplore and corresponding source code is deposited on GitHub.

**Author summary:**

- The mx-plore platform is a comprehensive database of miRNA:target interactions as described in the scientific literature.
- mx-plore is a new text-mining approach to extract context-dependent miRNA interactions from Pubmed abstracts and PMC full texts.
- mx-plore allows to query the comprehensive database in a context-specific manner, both on- and off-line.

## Introduction

Micro-RNAs (miRNAs) are small, non-protein coding RNAs that can post-transcriptionally regulate genes. The canonical mechanism works by binding of the respective miRNA to corresponding miRNA-binding sites on the regulated messenger RNA (mRNA), leading to a decay of the targeted mRNA [1], and, subsequently, less protein. Thus, miRNAs are powerful post-transcriptional regulators and allow targeted decay of mRNAs. miRNAs are also thought to be highly relevant in specific diseases, or even play an important role in orchestrating these [2]. It is well known that miRNAs work in context-dependent ways [3], meaning that it depends on the specific circumstances whether a miRNA is expressed, available, and possibly also whether it binds to certain non-canonical binding sites and leads to (fast) degradation.

There are several resources to explore the miRNA-gene-regulatory landscape known in many (model) organisms. More than 15 databases exist, which all collect miRNA-gene interactions (Table 1). However, only few of these databases consider contextual information for these interactions, such as mentioned diseases, processes, etc.

**Table 1.**
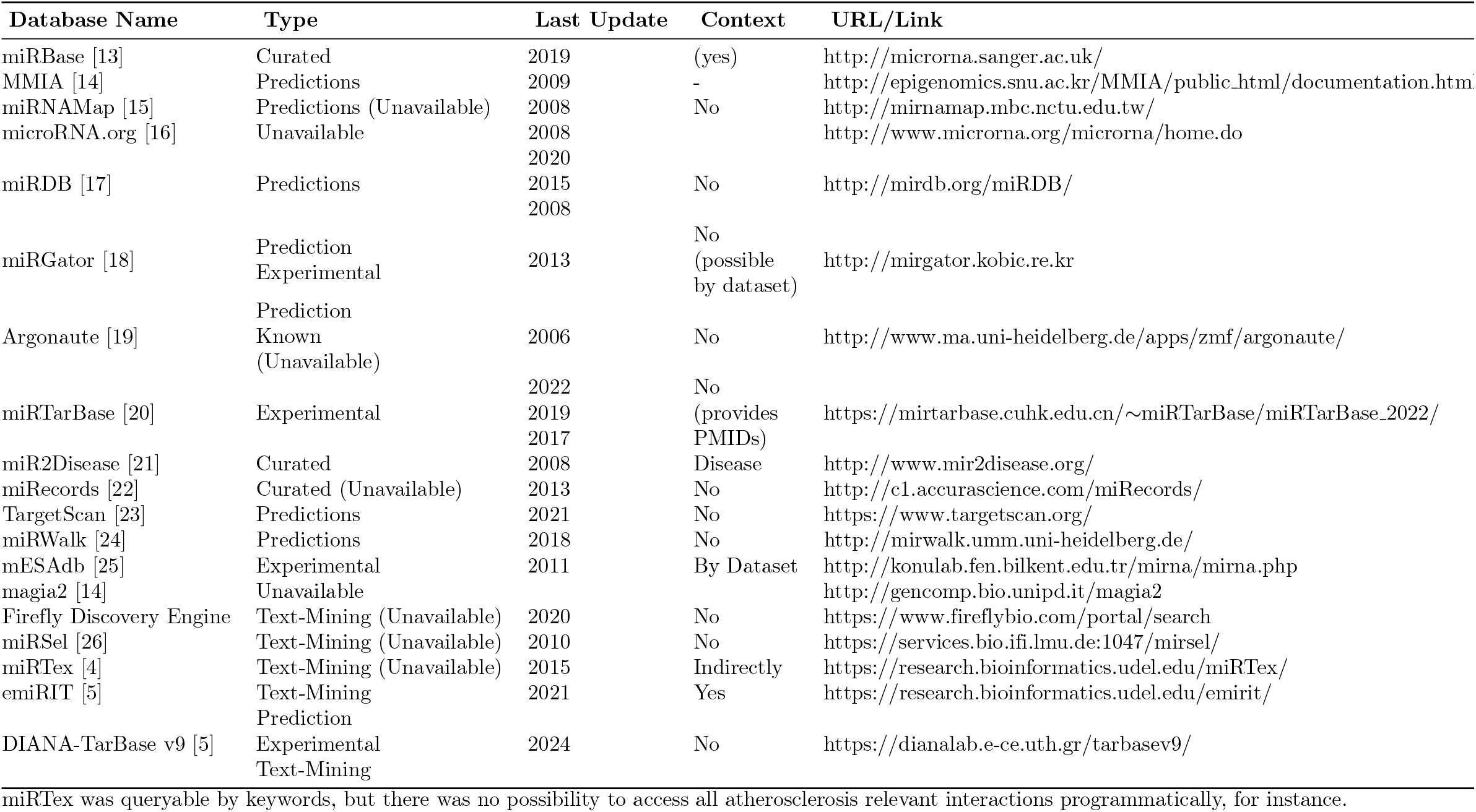
Existing databases for miRNA-gene interactions. Databases are annotated by type of origin of interactions, as well as their last update and whether they provide context information.

The two databases with such contextual information are miRTex [4] and emiRIT [5]. Besides pure text-mining resources, there are also experimental databases, such as DIANA-TarBase [6, 7] or miRTarBase [8, 9], which are regularly updated and have a long standing history of availability. This is, unfortunately, not true for most text mining resources. Moreover, text mining resources often only consider the abstracts of NCBI PubMed [10] in contrast to the larger text body of full text publications. Full texts provide more information, but are harder to interpret and analyse than abstracts, which are often written in more precise language [11].

Here we present a new rule-based method for miRNA-gene interaction mining that relies on pre-trained machine learning models for sentence dependency graph prediction. Comprehensible rules for both the retrieval of miRNA-gene interactions (mention-level prediction) and their putative regulatory direction are deduced and collected from both PubMed abstracts and PMC full texts into a database of miRNA-gene interactions. Our database is accessible online as a web platform, but can also be downloaded for programmatic access. For each evidence document of a miRNA-gene interaction, contextual information on identified cell types, biological processes and diseases is collected and displayed, which can further be used to filter for miRNA-gene interactions relevant to the user. This resource can serve as input for further integrative methods, including timelines, miRNA activity prediction and set-based over-representation analyses.

### Existing miRNA-gene databases

Several miRNA-gene-target-centered databases exist. A list of *some* databases is compiled in Table 1) These databases can generally be categorized into *manually curated, predictive* or *experimental* databases, based on their main content. Some formerly highly used databases have become unavailable, while others have emerged. Most of these databases lack context information, meaning they do not report the cell type, disease or process in which the miRNA-gene interaction was identified. One of the two databases with contextual information, miRTex [4], is several years old and is no longer accessible. When it was available, it did not allow systematic queries by contexts and it is not open source. The emiRIT database [5] integrates various resources of miRNA interactions, including miRTex. For each miRNA, its gene interactions are listed together with relevant process and disease information. However, with emiRIT, it is unclear on which text corpus the retrieval of miRNA-gene/disease interactions has been performed. The authors of miRTex state in their article that they analyzed both abstracts and full texts, but throughout their database description, they only refer to abstracts. For both miRTex and emiRIT, programmatic access or download is not possible. With regards to FAIR principles [12] and reproducibility, both resources lack essential parts.

## Materials and methods

### Accessing the corpus

Both Pubmed abstracts and PMC full texts (from the open access for commercial usage subset) were downloaded as JATS-formatted XML files in March 2024. Relevant information, such as authors and publication data, was extracted into tab-separated files. The entire abstracts and full text sections were taken from the XML-formatted files, and sentences were extracted using SpaCy with the tokenizer of the SciSpacy [27] model bionlp13cg (both version 0.2.4) for each input file.

### Named-entity recognition and extraction

The identification of gene and miRNA entities for which interactions can be later confirmed, or not is performed using named-entity recognition (NER) in mx-plore. The named entities are defined in a synonym file, which contains a synonym ID and multiple words that, if found, are associated with the respective synonym ID. For human genes, these synonyms are generated per human gene symbol using the HGNC annotation [28], previous gene symbols, and (previous) gene names. The same strategy is applied to synonyms for mouse genes using the MGI database [29]. For miRNAs, synonyms are created for all official human and mouse miRNAs extracted from miRBase [30]. In mx-plore, the focus is on the miRNA at precursor-level, meaning that -3p and -5p versions of a miRNA are not analysed separately. However, if such a distinction is desired, the entire mx-plore pipeline can be executed with a modified synonym list for miRNAs reflecting the desired detail level.

To find the named entities in text, a text mining approach similar to the one described by Hanisch et al. [31] is used. The *textmineDocument*.*py* tool is written in python3 and uses an Aho-Corasick data structure from the pyahocorasick library (https://github.com/WojciechMula/pyahocorasick) to find synonyms quickly and easily in text data. Because this approach is case-sensitive, a four-pass approach is used. The first pass detects the synonyms in a case-sensitive manner. The second pass uses a case-insensitive approach. The third and fourth passes employ the same strategies (case-sensitive and case-insensitive) but remove any white-spaces in sentence and synonyms. If multiple sentence files are given as input, the tool can also run them in separate processes in parallel. Searching through the entire set of PMC full texts takes about half an hour on 20 processes.

If a miRNA and a gene are found in the same sentence, both occurrences are checked for an interaction as described in the following sections.

### Extracting miRNA-gene interactions

The presented two step approach relies on accurate dependency graphs for the examined sentences. In order to derive such dependency graphs, the language processing framework SpaCy (v0.2.4) [32] is used. Best performance has been achieved using AllenAI’s SciSpaCy [27] sci-lg model (v0.2.4). AllenAI’s bionlp13cg model (v0.2.4) is used for term prediction. SpaCy predicts Part-of-Speech (POS) and dependency tags following the Universal Dependencies nomenclature [33].

### Finding miRNA-gene interactions

Using the following four rules on a sentence’s dependency graph prediction, it is determined whether a single miRNA and another gene entity form a miRNA-gene interaction (regardless of direction). The rules can be applied in any order, any only interactions which are confirmed through all rules are accepted.

### Conjugation Rule

The conjugation rule is used to determine whether two entities are in the same conjugation or not. Given a correct dependency graph, this is generally the case if the two entities (or any related words) are connected via a *conj* edge. For all elements of *conj* edges, all related elements must also be considered. Thus, for any word connected by *conj* edges, all connected dependencies through any of the following edges are followed and collected: *case, amod, nmod, dep, appos, acl, dobj, nummod, compound*.

However, some conjugations actually indicate an interaction and should not be rejected. For instance, conjugations like *we observed a direct regulation between miR-124a and Cxcr4* (Figure 2), should be kept. Hence conjugations with preceding *between* are retained.

**Figure 1.**
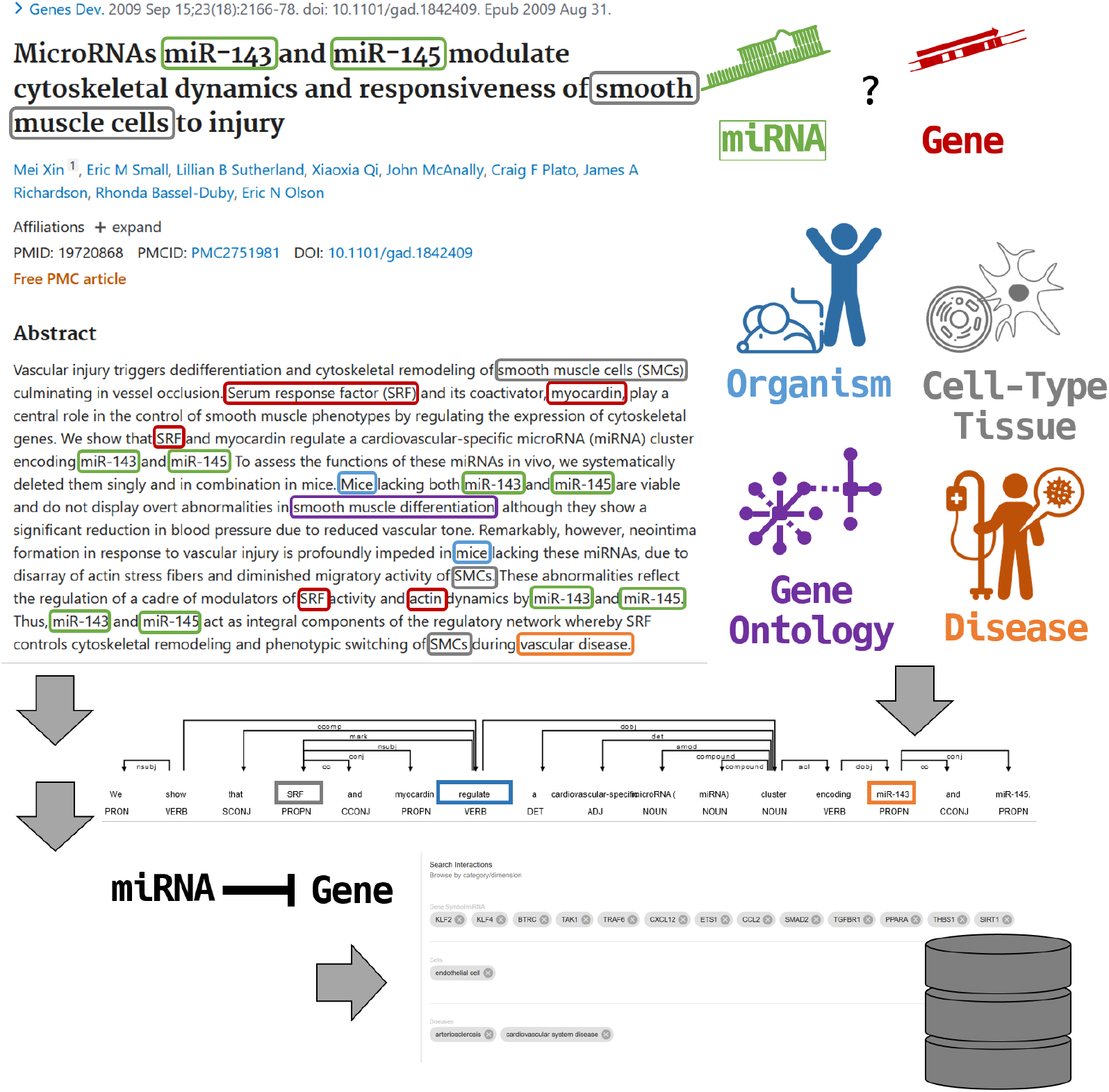
In the mx-plore system, full text publications and abstracts are used to identify relevant named entities. Based on that, miRNA-gene interactions are extracted, classified and stored in a database, which allows efficient querying.

**Figure 2.**
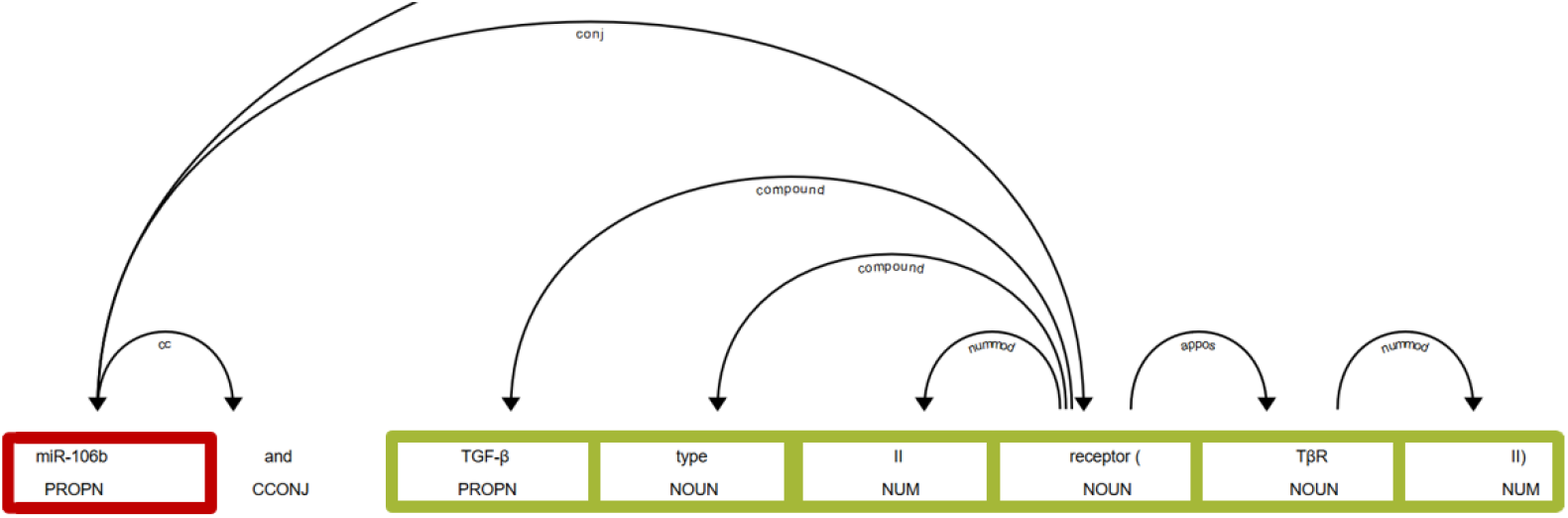
Conjugation rule example: *miR-106b* and *TGF-β type II receptor* are in the same conjugation, hence there is no interaction.

It was observed that there are many interactions mentioned between a miRNA or gene and a miRNA family or gene pathway. Thus, it is checked that the conjugation may not contain a *<miRNA*|*gene>* *pathway* -structure.

### SDP Rule

In previous work, a rule focusing on the intersection between the dependency paths from roots to the possibly interacting entities has already been applied [34]. Indeed, using the shortest path between two entities in the dependency graph has proven to be useful for general relation extraction [35] and has been applied to protein-protein interactions [36, 37]. Here, a similar concept is followed with the shortest dependency path (SDP). The SDP essentially contains words that connect two entities, thus capturing the most important concepts between these two entities.

On the SDP, it is verified that no subject is included unless it is one of the two observed entities. In the example (Figure 3), *miR-17/92* and *Shh* are not in a valid relation because *Shh* is a child of the actual subject *N-myc*. Therefore, the interaction is between *Shh* and *N-myc*, as well as *N-myc* and the miRNAs.

**Figure 3.**
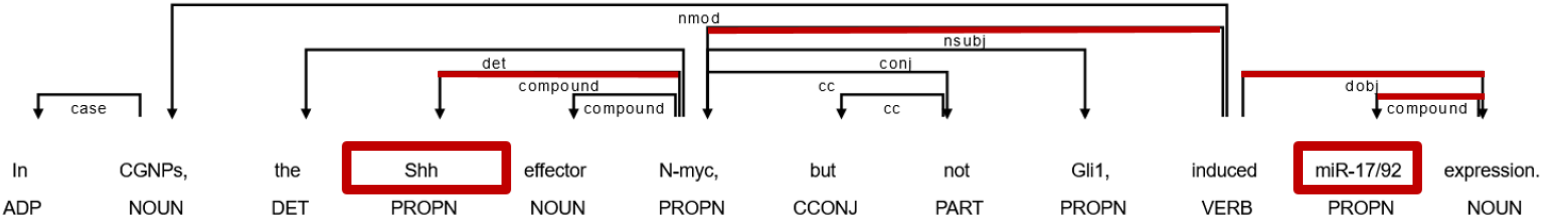
SDP rule example: *miR-17/92* and *Shh* are not in a valid relation, because *Shh* is a child of the actual subject *N-myc*. Hence the interaction here is between *Shh* and *N-myc*, and *N-myc* and the miRNAs.

Additionally, no *<verb> <noun> pathway* structure is allowed within the SDP to exclude pathway-related interactions.

### Compartment Rule

The compartment rule originates from the observation that interactions between miRNAs and genes typically occur within a sentence or clause, and are directly connected by a verb. For sentences consisting of multiple clauses, it is uncommon for a valid interaction to cross or skip such a subclause. Therefore, it is checked whether both entities are within the same subclause. To do this, the sentence is split into its subclauses, or compartments.

Given the following sentence, the four identified compartments are (listed directly below):

~~~
<e1>miR-200a</e1> was found to directly target beta-catenin mRNA,
thereby inhibiting its translation and blocking Wnt/<e2>beta-catenin</e2> signaling, which is frequently involved in cancer.
0 [miR-200a was found to directly target beta-catenin mRNA]
1 [thereby inhibiting its translation]
2 [blocking Wnt/beta-catenin signaling]
3 [which is frequently involved in cancer]
~~~

Before the actual check can be performed, all subclauses must be identified within the dependency graph.Utilizing the left-to-righ flow of the english language, the easiest way to extract these compartments is to split the sentence upon encountering special words or dependencies.

In general, a compartment is formed at one of the following dependencies: *cconj, xconj, conj, ccomp, parataxis, advcl, xcomp*. However, certain compartments are too fine-grained for further relation extraction. Therefore, depending on the observed token or dependency, further sub-rules are applied.

Compartments may be separated by several tokens, particularly their Part-of-Speech or dependency prediction ^1^. If a compartment is separated by a *VERB* (or *AUX*) tag, its full subtree will be further analysed. If the subtree starts with *because, through*, or other connecting verbs, it is not considered a separate compartment. The same applies to compartments starting with nouns. If the found *VERB* is connected by a conjugation, it is ensured that the full conjugation is within the compartment.

The compartment can also be split by the *amod* dependency. For an *amod* dependency, the sentence must be split by *thereby, suggestive, while*, etc.

A *VERB* with an *acl:relcl* dependency must also have split-words like *whereby* at the start of the subtree.

Finally, a compartment can also be split by a *conj* if it starts with a splitting word like *actually*.

Some compartments are not divided by verbs or conjugations. Therefore, compartments are also split at any occurrence of a semi-colon (;).

The final compartment check ensures that both miRNA and gene are contained within the same compartment.

### Context Rule

There exist several words which make a miRNA or gene entity not a target entity for the desired interactions. This rule is the most heuristic one in this framework and may need to be adapted for other use cases, such as identifying protein-protein interactions. For instance, consider the previously discussed sentence:

~~~
<e1>miR-200a</e1> was found to directly target beta-catenin mRNA,
thereby inhibiting its translation and blocking Wnt/<e2>beta-catenin</e2 signaling, which is frequently involved in cancer.
~~~

The interaction (*e*1, *e*2) in this example sentence is not about the miRNA influencing the *Wnt* or *beta-catenin* genes, but rather their related signaling pathways. Therefore, (*e*1, *e*2) does not describe a direct miRNA-gene interaction.

In the context rule, the text before and after a miRNA or gene entity is checked for the presence of certain keywords. If any one of these keywords are found, the interaction is rejected. This includes, for example, checks in order to determine whether a gene-entity is related to a specific cell- or genotype, or whether miRNA-gene complexes are formed. To achieve this, the closure around the gene entity is considered.

### Determining miRNA-gene regulation

After finding a miRNA-gene interaction, this interaction is meant to be further characterized. The direction of the interaction (be it miRNA regulates gene, or vice versa) as well as the direction of the regulation (up-/down-regulated) is of interest.

In order to determine both interaction properties, again a rule-based approach is chosen. For determining relations it was heavily relied on the dependency graph prediction (see previous section). However, it has been observed that this prediction is incorrect in many details. These details, however, are essential for solving the problem of determining in which direction a regulation occurs.

Hence it was decided to not use the dependency graph as major analytical object of interest. Instead, this approach heavily relies on one property of the English language: the scrambling of words is uncommon. While many languages, such as German for instance, make heavy use of scrambling, in English the use of a pragmatic word order is much more tightened [38]. Hence most sentences follow the Subject-Verb-Object scheme, also in this order. Also words, which associate with the start of a sentence, are usually found at the beginning of the sentence.

The following rules hence operate on the stems of certain word-groups, such as the stem *up-regul* for any word in this family, like *up-regulating* or *up-regulator*. These stems have been collected from the training data on the one side, and manual curation from reading literature on the other side.

The following rules build upon each other and are applied one after another. This means, that only if a rule delivers a prediction for the miRNA-gene interaction in a specific sentence, no further rule is checked. The rules are applied in the order of their mention in this manuscript to the miRNA-gene interaction in a sentence.

### Compartment and Between Rule

Both the compartment and between rule check for the presence of specific context descriptions between the corresponding miRNA and gene words within a sentence. In the given example sentence (Figure 4), first the compartment containing miRNA and gene is determined, which is defined as both entities and all words in-between. For the compartment rules, it is searched for specific keywords, such as *negatively correlates* or *positive correlation*, within the compartment. Depending on the found keywords and the order of miRNA and gene entity, the interaction direction is determined (here MIR GENE) as well as the direction of the regulation (DOWN). The *Between* rule works similarly, but accepts a broader set of terms (e.g. such as *recognizing* or *binding efficiency*) and enlarges the considered text also by word tokens attributed to the miRNA and gene entities via the dependency graph for positive and negative interactions. Neutral interactions must be detected within the narrowed compartment.

**Figure 4.**
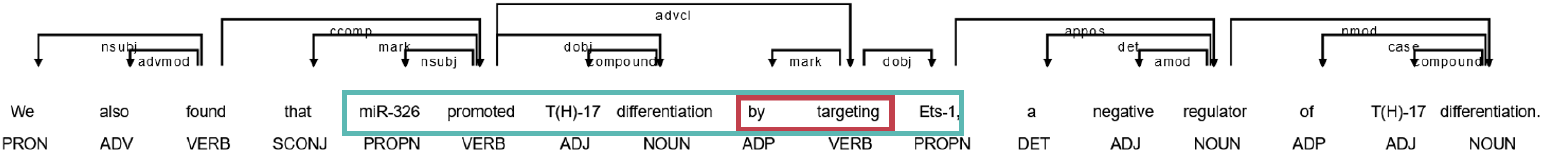
Interaction Compartment Rule.

### Counts Rule

If no direct associations are found, the focus has to be led on a broader stem-based approach, using a list of pre-defined word stems with corresponding directional annotation (e.g. positive, negative or neutral). For both the miRNA and the gene, any stem within the proximity of the word is identified, and its direction is counted. Neutral stems are ignored at this stage. In the example given in Figure 5, the searched proximity is marked in dark-green. For the miRNA one negatively associated stem is found (inhibition) and for the gene a positively associated one (increase). Then it is checked whether the miRNA and gene directions are inverted, that means miRNA negative, and gene positive, or vice versa - which is the case here. If this is the case, a MIR GENE DOWN regulation is predicted. Otherwise the *counts between* rule is considered, which counts the number of detected stems for each entity and assigns the interaction direction accordingly. If the number of detected positive stems equals the number of detected negative stems, a neutral interaction is predicted.

**Figure 5.**
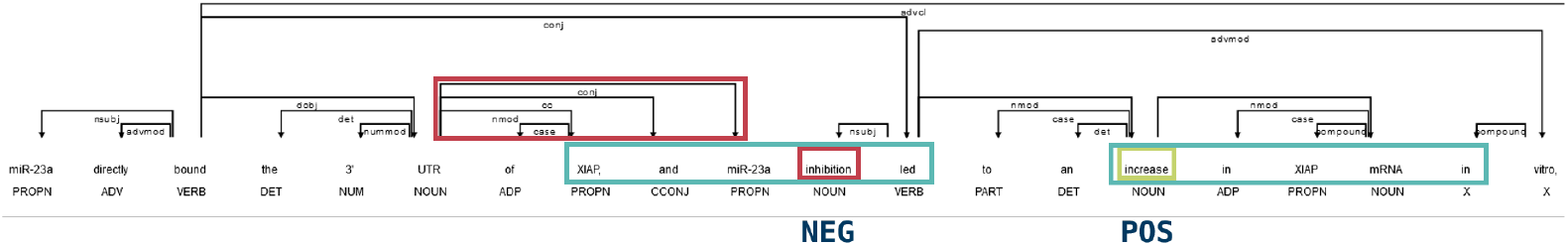
Interaction Counts Rule.

### Return Rule

The final rule has to care of any yet unassigned interaction. For this rule, all stems which are between both entities are considered. The most-frequently occurring stem direction is predicted. The interaction direction is determined by the order of the entities, and whether passive is detected or not. In the example given in Figure 6, SRF stands before miR-143 in a non-passive sentence. Hence the interaction direction is GENE MIR. The found stem, *regulate* is neutral (not knowing up- or down-regulating). Hence the predicted interaction is a GENE MIR NEUtral interaction.

**Figure 6.**
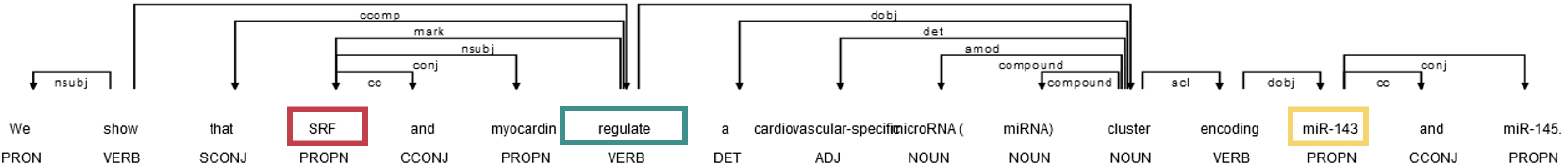
Interaction Final Rule.

### Benchmark

For evaluating both the interaction and regulatory prediction methods, the benchmark provided by Bagewadi et al. [39] is used. This benchmark allows to compare the proposed method also with other, already published, methods, like miRTex [4], as these also used this benchmark for comparison. The benchmark consists of a training dataset (201 documents, 397 interactions) and a test dataset (100 documents, 232 interactions). For 24 interactions (see Supplemental Material) the gold standard was changed to reflect a common handling of miRNA-pathway interactions (see Context Rule above).

The original benchmark has been extended, as part of this work, to also include the interaction type (miRNA → gene, gene → miRNA) and regulatory direction (down, up, neutral) for all miRNA-gene interactions of the original benchmark.

### Web-Portal

The database can be accessed in two ways. Through mx-plore it is possible to query the miRNA-gene interactions interactively and without programming skills. The website is provided using flask and sqlalchemy for accessing the sqlite database. The actual frontend is implemented in HTML and javascript, making use of bootstrap, data tables, jquery and yadcf. This flask application not only serves the frontend but also provides routes to access detailed information or the timeline plot. The second way is to download the sqlite database from the website, such that the results can also be accessed programatically.

On the web portal and on the detail pages for each interaction, several information a displayed: First the interaction itself is shown (e.g. miRNA targets gene, or miRNA down-regulates gene). Next, the interaction context is shown in terms of identified gene ontology, cell line and disease ontology terms identified in documents in which the specific miRNA-gene interaction has been identified. This is followed by the gene structure of the given gene as well as predicted (conserved) binding sites as contained in the TargetScan v8.0 human or mouse data [23] using the javascript version of IGV [40]. Below these information, all sentences used as evidence for the selected miRNA-gene interaction are shown and linked to the respective abstracts and documents. Finally a timeline plot, generated using d3-milestones (https://github.com/walterra/d3-milestones) is shown, setting the evidence documents into a temporal context.

### Timelines

For a specific miRNA-gene interaction all literature evidence can be collected. Using publication date and contextual information, these can be displayed on a time axis where the x-axis location corresponds to the publication date. Single publications are annotated with each the two most common diseases, GO terms and cell types/lines. In order to determine the most common terms per ontology, for each term identified for the document, a count of 1 is added. All respective parents are added with a count of 2, in order to select informative, yet not too specific terms. Finally, the terms with the highest scores are chosen to be displayed.

### Data Availability

The whole interaction retrieval pipeline is published online with source code and required steps for reproducing the results (or updating them). All source code is available from GitHub https://github.com/mjoppich/mxplore. A corresponding conda environment with execution instructions is available. The database is accessible from https://rest.bio.ifi.lmu.de/mxplore, where it is also offered for download. The text mining results will be deposited at the time of publication at Zenodo (https://doi.org/10.5281/zenodo.12704497).

## Results and Discussion

mx-plore is an integrative text mining framework for miRNA-gene interactions. The mx-plore framework consists of multiple parts, as shown in Figure 1. From PubMed abstracts and PMC full texts entities are extracted using named-entity recognition (NER). From these, miRNA-gene interactions are identified using the structure of sentences in which a miRNA-gene co-occurrence was identified. Together with contextual information of the relevant document (e.g. organism, cell type, gene ontology or diseases), the miRNA-gene interaction and its regulatory direction is saved in a database and made available for query from the web, or as download.

### Evaluating miRNA-gene mention detection

The miRNA-gene interaction detection is a detection on the mention level (without any direction information). That means, that in this step it is decided whether two entities form a valid miRNA-gene interaction or not.

In a previous study, atheMir [34], we found that our NER works considerably well in conjunction with a rule-based approach, and performs as good as or even better than field experts do - also in a context sensitive way. For that approach we used a three-rule approach: for a valid interaction between two entities, these must not be within the same conjugation, be connected by a verb and share at least one element of the paths from the root to the respective entities in the dependency graph. However, in contrast to other existing techniques and methods, this 3-rule based approach could be improved.

Using the training data set from the benchmark [39], the four rules presented in Section Finding miRNA-gene interactions have been developed. The performance of mx-plore and each combination of rules has been evaluated using the 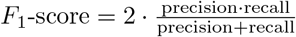 (Figure 7, Table S2). No rule (always accept) performs worst (in terms of the *F*_1_-score), followed by the single rules (only one rule applied). It is interesting to note that the SDP-only rule performs worst of all, and worse than the other single rules. The single rules are followed by the double-rules which are followed by the triple-rules. Finally the use of all available rules together delivers the best results. Hence all rules are applicable and there is no single rule which outperforms all others. This also suggests that the *problems* provided by the benchmark are considerably different.

**Figure 7.**
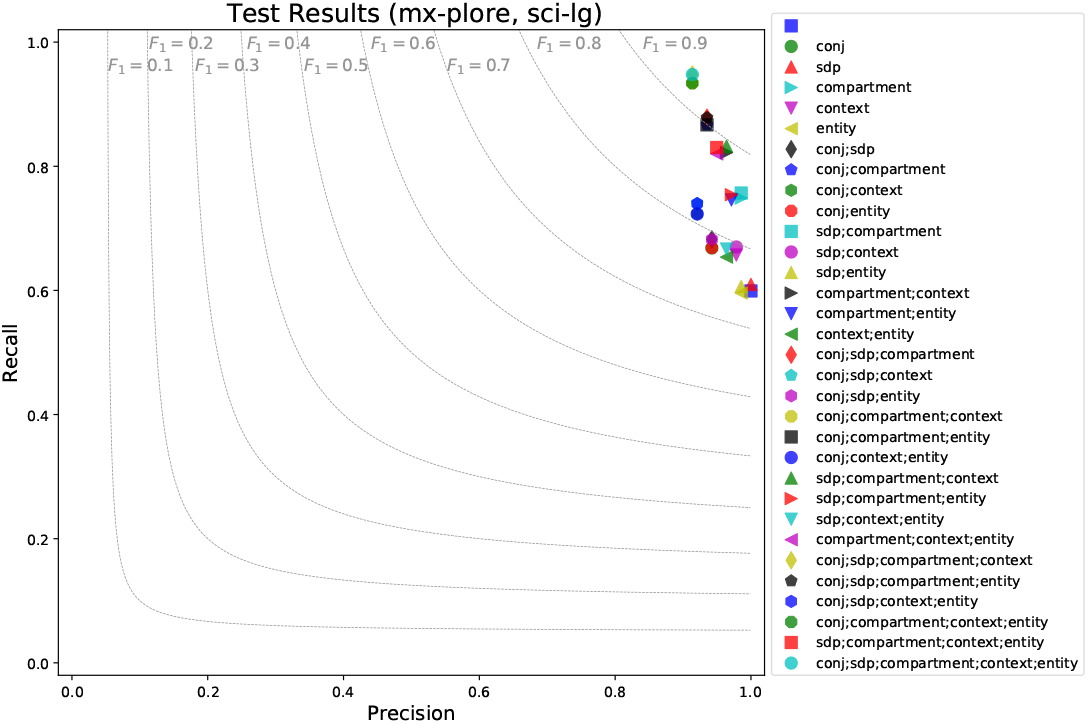
mx-plore performance evaluation using sci-spacy (large) model. For each rule combination the mx-plore predictions are evaluated on the benchmark’s test dataset. It can be seen that with more rules, results become better in both precision and recall.

### It depends on the dependency prediction

During the interaction extraction phase, mx-plore makes particular use of the dependency graph generated by spacy [32]. All applied rules rely on a correct dependency prediction, particularly the shortest dependency path (SDP) rule. While the other rules also rely on the dependency graph for recognizing conjugations or compartments, the acceptance in the SDP rule depends directly on the dependency graph. Hence the question arises how much the success of the rule-based method depends on the dependency prediction.

In the regular configuration mx-plore uses scispacy’s large model for dependency prediction. But there is also a special model trained on the biomedical BIONLP13CG corpus. Likewise, spacy also comes with a regular large model, trained on general text, like newspapers or *trashy* literature. These additional models were also evaluated on the test dataset (Figures S1, S2). The general observations made for the scispacy large model remain valid for the other models: the more rules are added/applied, the better the predictions get. However, one specific observation for the SDP rule in the general spacy-model is interesting: Here the SDP rule performs worse than no rule, and even the difference between the 3-rules and all rules becomes small. This is however not surprising: in the previous iteration of this text mining framework, atheMir [34], it was noticed that several incorrect interactions have been reported due to incorrectly resolved dependency paths to the root, particularly around biomedical words. Hence, using a model not trained for such entities, many incorrect SDPs will lead to a result even worse than the no-rule concept.

Looking at the absolute *F*_1_ values it can be noticed that on choosing the wrong model and applying the same rules, the score varies from 0.95 (scispacy large) down to 0.77 with the spacy large model. This highlights the major impact the dependency prediction has on even a rule-based interaction detection approach. But moreover this highlights the need for well and correctly pre-trained models. Considering that the rule-based approach here resembles a fine-tuning of a pre-trained deep learning model, it can be argued that a badly pre-trained main model (e.g. pre-trained on different kind of text, not biomedical) can not be rescued by a good fine-tuning.

### Comparing all methods

While it could be seen that all four rules combined perform considerably well, the key question is to analyse how well this rule-based approach performs - compared to other approaches and resources. With the chosen benchmark this comparison can be performed: the authors of the benchmark already compare multiple techniques on this benchmark, and the miRTex authors [4] rebuilt the miRSel approach and present their own approach, also evaluated on this dataset. All these results, together with the ones discussed so far, have been combined and are presented in Figure 8 and Table S1. Results marked with † are taken from [39] and results marked with ‡ are taken from [4].

**Figure 8.**
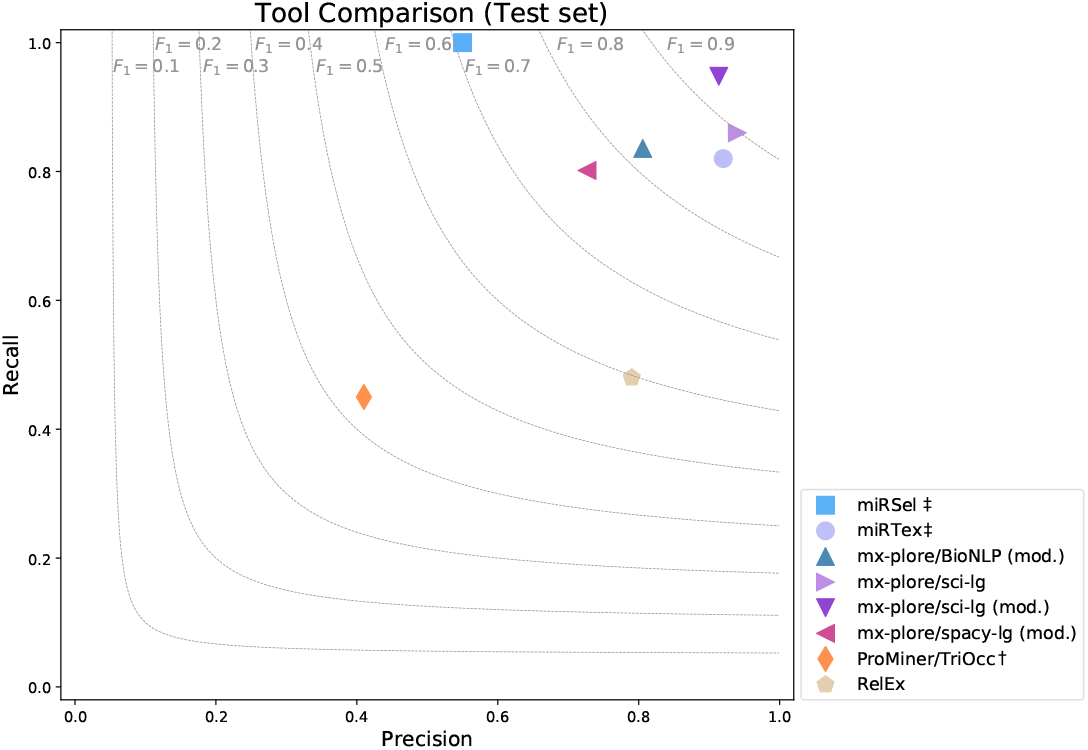
mx-plore performance in comparison with other tools. It can be noted that all tools perform better than a simple tri/co-occurrence approaches (ProMiner). But also mx-plore improves in contrast to its previous version, atheMir. This holds for any spacy model employed. This can be explained by the usage of more suitable rules. miRSel and RelEx perform well, but even range behind mx-plore. Particularly RelEx has problems with specific sentences, probably due to an incorrect dependency resolution. Finally mx-plore using the scispacy large model achieves the top *F*_1_ score of over .9.

A simple co-occurrence based analysis achieves a *F*_1_ score of less than 0.5 [39], followed by the (general) relation extraction tool RelEx [41]. The re-implementation of miRSel achieves a high recall with low precision, and the three-rule-based atheMir performs slightly better with a *F*_1_ score of 0.75. miRTex outperforms mx-plore on the regular spacy model as well as the smaller BIONLP model - only slightly on the latter. However, the large scispacy model combined with the rule-based approach of mx-plore outperforms even miRTex.

Even though miRTex is also rule-based, one design decision of miRTex, which it has common with RelEx, might be a reason for the reduced recall: no trigger word should be needed for a valid miRNA-gene interaction. There are virtually so many suitable trigger words used in literature, that it is simply not possible to find and enumerate them all. Instead, extensive use of the dependency graph should be made. While the dependency graph has problems of its own (and hence is not be used for determining the direction of regulation), it is good enough to identify whether two entities are in some kind of relation or not. Particularly the dependency graph can be used to reject entities which occur together in a conjugation. That said, having a correct dependency graph is crucial. Modern dependency predictions have become quite reliable, even on unknown sentences and words. Being trained on actual biomedical literature improves their performance such that it can be used reliably to derive interactions.

### Evaluating miRNA-gene regulation detection

Within mx-plore, a two step approach is chosen: First an interaction is detected, then the interaction direction. Using the extended benchmark, the four rules for predicting the interaction direction of an identified interaction presented in Section Determining miRNA-gene regulation were designed according to the training data. These rules make use of two observations: (1) while the dependency tree is able to identify whether miRNA and gene are not interacting, it is too coarse to identify the direction of a regulation. (2) scrambling of words is very uncommon in the English language [38], hence determining the direction of the interaction using word stems, not relying on the dependency graph, is possible. On a side note: the red marked dependencies in Figure 5 show that the dependency prediction is sometimes problematic. For instance the conjugation around the miRNA is not resolved correctly. While this does not hamper the interaction detection here (still same compartment, not same conjugation, just longer SDP), the interaction direction detection could be influenced, because there is no relation between inhibition and miR-23a. This shows, exemplarily, why for this task the stem-based approach is more suitable.

Again the single rules and any combination of rules has been compared (Figure 9). For the no-rule case it is always predicted that the miRNA regulates the gene down. Some rules perform even worse than this simple rule. This, however, is explainable at the example of the return rule. This rule is designed to be used only in specific cases, and after other cases have already been handled by previously checked/applied rules. Using this rule right-away is not a representative use-case. In contrast to the interaction detection, here the rules are incremental and cannot be seen individually. Nonetheless, with more added rules, an improved *F*_1_ score can be observed. Again, all rules taken together perform best with a *F*_1_ score of 0.88. However, in general, with more rules, prediction results improve.

**Figure 9.**
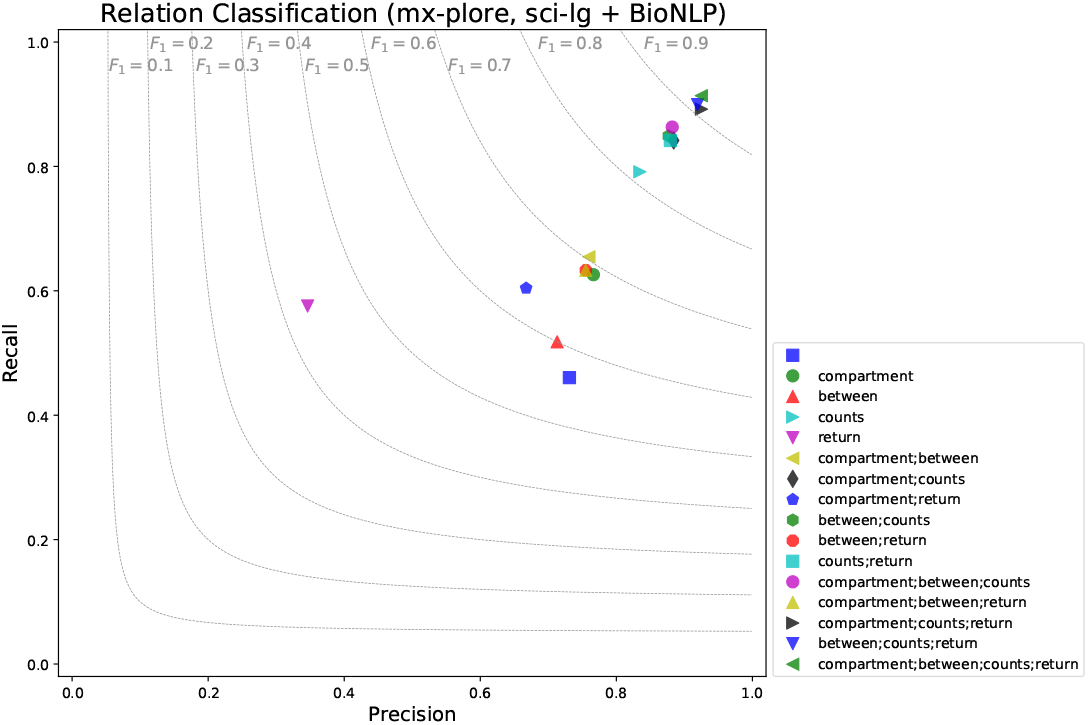
mx-plore interaction direction evaluation. The extended benchmark is used to check the prediction miRNA-gene interaction and direction. While the previous checks focused on detecting an interaction, here it is checked which entity regulates the other entity into which direction. Using no rule (and always predicting MIR GENE, NEU), performs quite well. Some rules, taken out of sequence, perform even worse.

### Database of miRNA-gene interactions from text mining

Using the mx-plore pipeline, many miRNA-gene interactions could be discovered. An overview of all found interactions, their type and their source is given in Table 2. mx-plore provides a web-based user interface for quick and easy access. The user can enter specific genes or miRNAs of interest. Also the user can select which context the returned interactions must be found in. Therefore the user can enter terms from all ontology terms in the specific dimensions. Returned search results are subset to only those miRNA-gene interactions which are found in literature associated with the search word. For programmatic access, mx-plore offers to download the database in sqlite format (including annotations), as well as the interactions as tab-separated file.

**Table 2.**
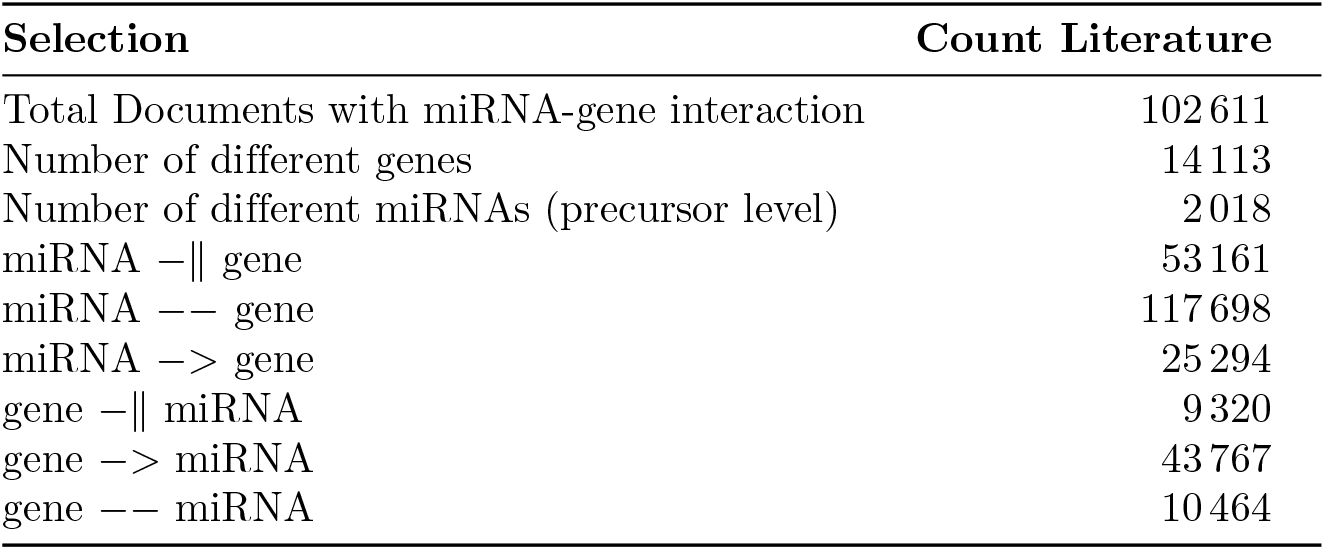
Overview of text mining-based miRNA-gene interactions. Certain interactions might, for example, be recorded as both neutral and repressing regulation and hence are counted twice. Interactions are designated with −∥ for a DOWN-regulation, −− for NEUtral interaction and −*>* for UP-regulation.

When researching about a specific miRNA-gene interaction it is interesting to know in which *context* this interaction has already been researched on. Using the *Timelines* plot, it is possible to plot in which context the specific miRNA-gene interaction has already been looked into, and when. Results from mx-plore are taken, condensed, and plotted. In the shown example of a miRNA −*>* gene interaction (Fig. 10) between miR-222 and CXCL12 it can be seen that there are 7 evidence documents and that this interaction is the consensus interaction, meaning the most prevalent interaction between miRNA and gene. Below this general information the contextual keywords identified from the evidence documents are displayed in a word cloud. The more frequent a context term is, the bigger it is displayed. Here, it can be seen that this interaction plays an important role in angiogenesis and apoptosis. Next follow the gene structure of the studied gene (CXCL12) as well as predicted miRNA binding sites, taken from TargetScan version 8 [23]. After this the interaction evidences, namely all occurrences of the reported miR-222 and CXCL12 interaction in texts, are displayed with the miRNA and gene being highlighted, and links to the respective articles given. The detail page closes with a timeline plot for the miRNA-gene interaction showing on the one hand the temporal progression on research about the specific interaction, and on the other hand showing in which process-, cellular- and disease-context and by whom the respective interaction has been studied. This allows users to directly judge whether the selected miRNA-gene interaction is of direct interest.

**Figure 10.**
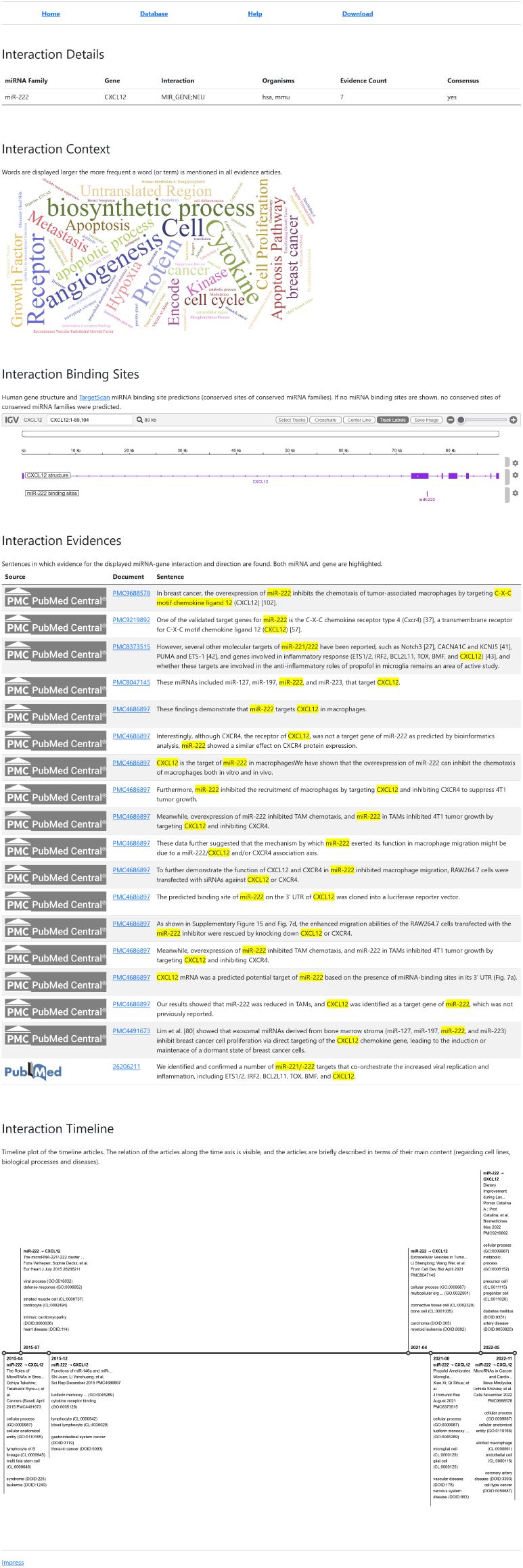
Detail page of the mx-plore web portal. The detail page for a selected miRNA-gene interaction. For the selected miRNA miR-222 and the gene CXCL12 and the direction miRNA −∥ gene, first the number of evidences is listed along the boolean flag whether this is the consensus interaction direction. Below the summary a text cloud with the mentioned context information for this interaction is shown, followed by the actual evidences and links to the respective documents. At the bottom a timeline plot is showing when research on the specific interaction was conducted and in which disease-, cellular- and process-context.

Since it has been noticed that existing databases vanished suddenly in the past, attention has been paid on making this resource endure for a long time. In terms of FAIR principles this is of highest importance, because only an accessible database, with resources and methods published, ensures that information can be looked up even after several years. Hence, not only a web portal with access to the database is available, but also the source code, instructions for running the text mining and post-processing for receiving all miRNA-gene interaction as well as a dump of the whole database is made available. In the light of the reported half life of bioinformatics web-services of 10 years [42] this ensures persistent access to data.

## Conclusion

In this paper the miRNA-gene interaction database mx-plore is described. The interactions were retrieved from text mining PubMed abstracts and PMC full texts. The miRNA-gene interaction extraction has been benchmarked on a public benchmark and could hence be set into context with existing resources. The mx-plore interaction mining outperforms all existing methods,. However, it becomes apparent that even rule-based interaction mining platforms require accurate dependency graph prediction. When using model-based methods for dependency graph creation, it is important to use a model trained on the same origin of text: biomedical literature.

With the provided mx-plore web portal it is possible to easily search miRNA-gene interactions within literature, and to filter results such that only interactions also found in specific process-, cell type- and disease-contexts are shown. The context of the miRNA-gene interaction is also visualized using a word cloud on the interaction detail page. On this page also all text evidence for that miRNA-gene interaction are listed such that the user can easily see whether the interaction is of relevance, or not. While it was shown that the interaction detection as well as the regulation prediction work accurately, these are no fail-proof processes. With the evidence sentences shown on the detail page and the timeline, the user can also judge whether to trust the interaction. Finally the timeline sets all work already performed on a miRNA-gene interaction into context and allows the user to dig deeper into the topic by reading the most recent, or context-wise, most relevant paper.

mx-plore is available online as web portal, but can also be downloaded as aggregated sqlite database with all context annotations, evidence sentences and publication information. This enables users to also use the database programatically in custom methods and analyses.

With this resource we hope to contribute to this highly innovative field, enabling scientists to verify miRNA activity, and to relate own findings with existing knowledge.

## Supporting information

**Figure S1.**
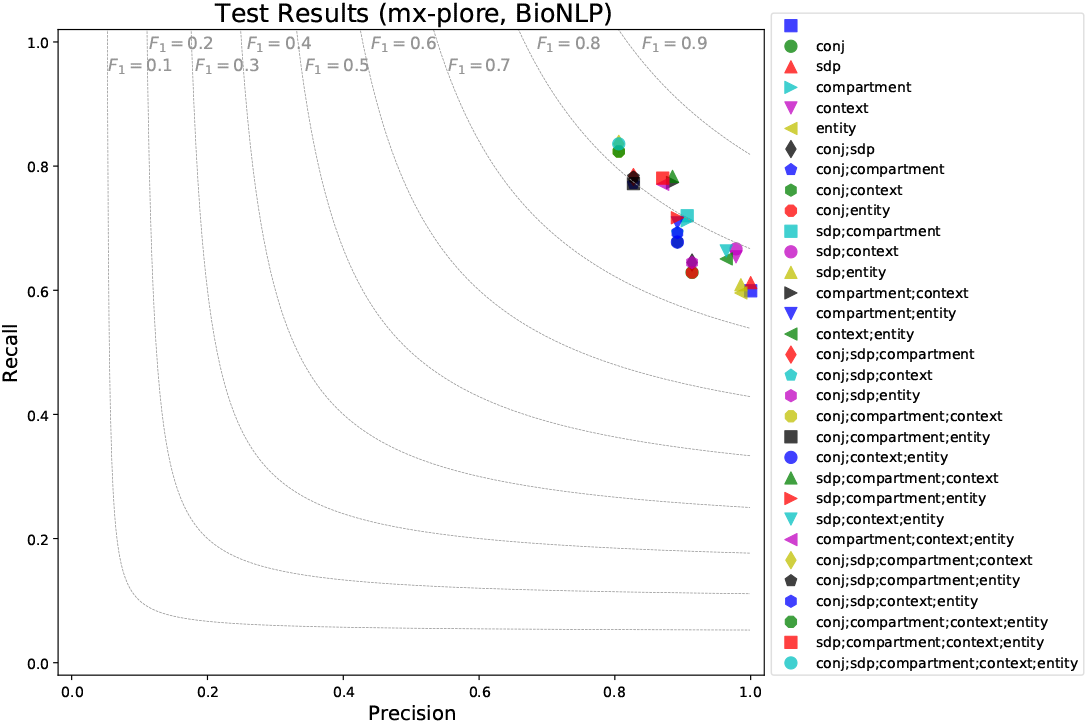
Detailed mx-plore performance for miRNA-gene interaction prediction using the sci-spacy (BioNLP) model on the test data.

**Figure S2.**
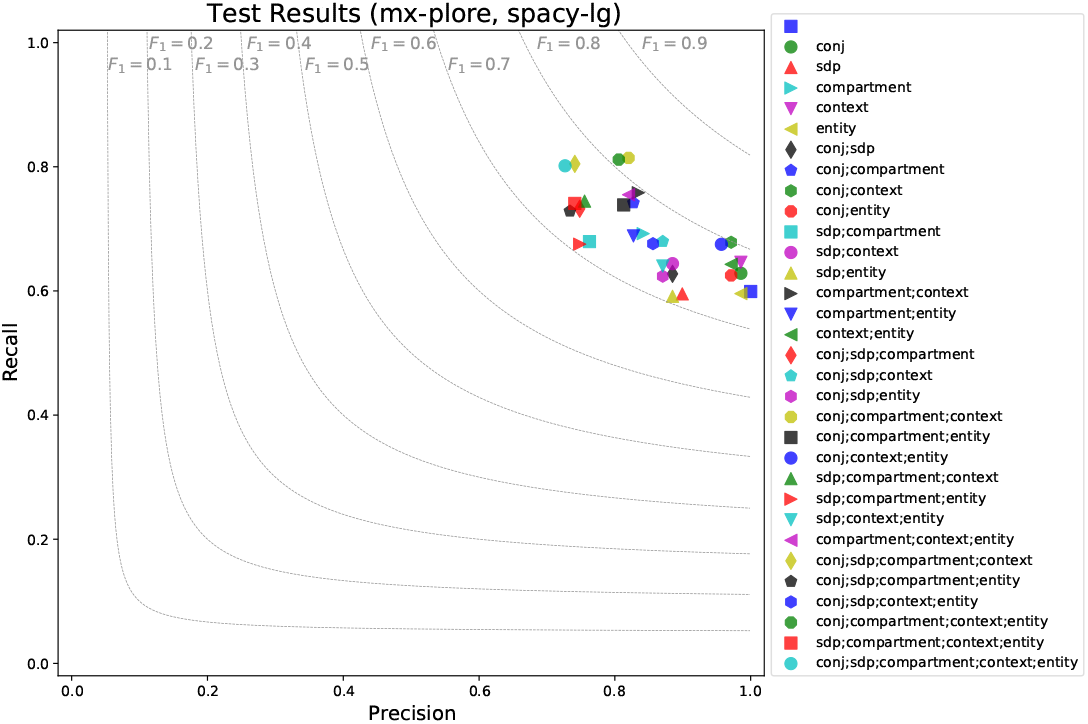
mx-plore performance evaluation using spacy (large) model. In contrast to the evaluation in Figure 7, here the default spacy (large) model was used for dependency graph prediction. It can be seen that the results are worse than for the specific sci-spacy model. Moreover, the SDP-only rule here performs even worse than the *always accept* rule, at least in terms of *F*_1_ score.

**Table S1.** Performance comparison between mx-plore and other tools. Results designated by † are taken from [39] and those with ‡ are taken from [4]. Results designated with (mod.) are evaluated on the modified test data. All other results are evaluated on the unchanged test data from [39].

| Rules enabled | Precision | Recall | $F_1$ |
| --- | --- | --- | --- |
| miRSel ‡ | 0.550 | 1.000 | 0.710 |
| miRTex ‡ | 0.920 | 0.820 | 0.867 |
| mx-plore/BioNLP (mod.) | 0.806 | 0.836 | 0.821 |
| mx-plore/sci-lg | 0.940 | 0.860 | 0.898 |
| mx-plore/sci-lg (mod.) | 0.914 | 0.948 | 0.930 |
| mx-plore/spacy-lg (mod.) | 0.727 | 0.802 | 0.762 |
| ProMiner/TriOcc † | 0.410 | 0.450 | 0.429 |
| RelEx | 0.790 | 0.480 | 0.597 |

**Table S2.**
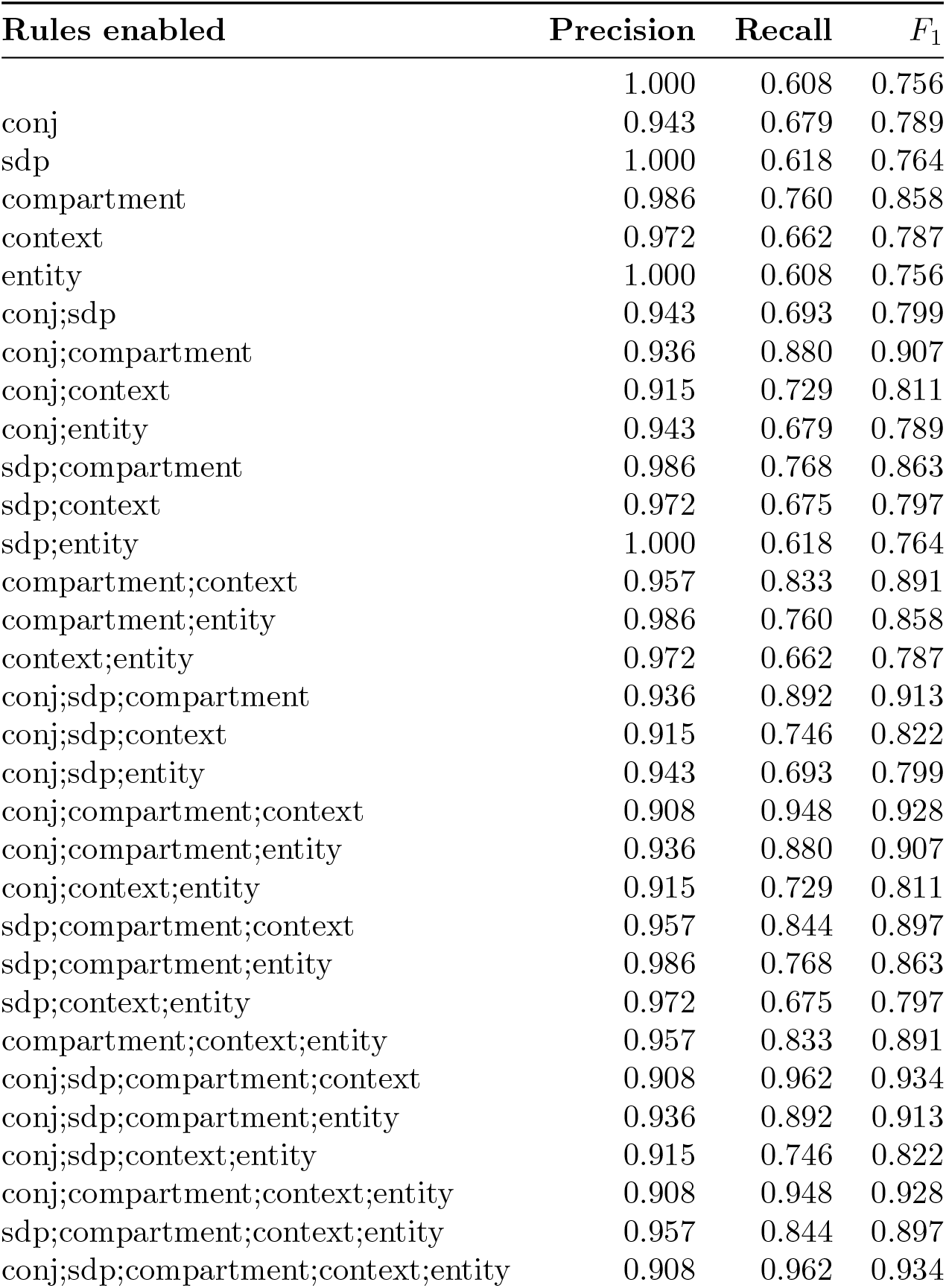
mx-plore performance evaluation using sci-spacy (large) model.

| Rules enabled | Precision | Recall | $F_1$ |
| --- | --- | --- | --- |
|  | 1.000 | 0.608 | 0.756 |
| conj | 0.943 | 0.679 | 0.789 |
| sdp | 1.000 | 0.618 | 0.764 |
| compartment | 0.986 | 0.760 | 0.858 |
| context | 0.972 | 0.662 | 0.787 |
| entity | 1.000 | 0.608 | 0.756 |
| conj;sdp | 0.943 | 0.693 | 0.799 |
| conj;compartment | 0.936 | 0.880 | 0.907 |
| conj;context | 0.915 | 0.729 | 0.811 |
| conj;entity | 0.943 | 0.679 | 0.789 |
| sdp;compartment | 0.986 | 0.768 | 0.863 |
| sdp;context | 0.972 | 0.675 | 0.797 |
| sdp;entity | 1.000 | 0.618 | 0.764 |
| compartment;context | 0.957 | 0.833 | 0.891 |
| compartment;entity | 0.986 | 0.760 | 0.858 |
| context;entity | 0.972 | 0.662 | 0.787 |
| conj;sdp;compartment | 0.936 | 0.892 | 0.913 |
| conj;sdp;context | 0.915 | 0.746 | 0.822 |
| conj;sdp;entity | 0.943 | 0.693 | 0.799 |
| conj;compartment;context | 0.908 | 0.948 | 0.928 |
| conj;compartment;entity | 0.936 | 0.880 | 0.907 |
| conj;context;entity | 0.915 | 0.729 | 0.811 |
| sdp;compartment;context | 0.957 | 0.844 | 0.897 |
| sdp;compartment;entity | 0.986 | 0.768 | 0.863 |
| sdp;context;entity | 0.972 | 0.675 | 0.797 |
| compartment;context;entity | 0.957 | 0.833 | 0.891 |
| conj;sdp;compartment;context | 0.908 | 0.962 | 0.934 |
| conj;sdp;compartment;entity | 0.936 | 0.892 | 0.913 |
| conj;sdp;context;entity | 0.915 | 0.746 | 0.822 |
| conj;compartment;context;entity | 0.908 | 0.948 | 0.928 |
| sdp;compartment;context;entity | 0.957 | 0.844 | 0.897 |
| conj;sdp;compartment;context;entity | 0.908 | 0.962 | 0.934 |

**Table S3.** mx-plore interaction direction evaluation.

| Rules enabled | Precision | Recall | $F_1$ |
| --- | --- | --- | --- |
|  | 0.732 | 0.454 | 0.560 |
| compartment | 0.766 | 0.617 | 0.683 |
| between | 0.713 | 0.511 | 0.595 |
| counts | 0.837 | 0.794 | 0.815 |
| return | 0.353 | 0.582 | 0.439 |
| compartment;between | 0.759 | 0.645 | 0.697 |
| compartment;counts | 0.886 | 0.844 | 0.864 |
| compartment;return | 0.670 | 0.610 | 0.639 |
| between;counts | 0.878 | 0.851 | 0.865 |
| between;return | 0.757 | 0.638 | 0.692 |
| counts;return | 0.881 | 0.844 | 0.862 |
| compartment;between;counts | 0.884 | 0.865 | 0.874 |
| compartment;between;return | 0.757 | 0.638 | 0.692 |
| compartment;counts;return | 0.926 | 0.894 | 0.910 |
| between;counts;return | 0.921 | 0.901 | 0.911 |
| compartment;between;counts;return | 0.926 | 0.915 | 0.920 |

## Author Contributions

MJ: Conceptualization, Data Curation, Formal Analysis, Investigation, Methodology, Software, Validation, Visualization, Writing – Original Draft Preparation, Writing – Review & Editing; SK: Software, Writing – Review & Editing; RZ: Conceptualization, Resources, Funding Acquisition

## Footnotes

1 https://spacy.io/usage/linguistic-features/

